# Intrinsic apoptosis is evolutionarily divergent among metazoans

**DOI:** 10.1101/2021.12.21.473695

**Authors:** Gabriel Krasovec, Éric Quéinnec, Jean-Philippe Chambon

## Abstract

Apoptosis is regulated cell death that depends on caspases. Upstream of each apoptotic signalling pathway is involved a specific initiator caspase. Characterised in nematode, fly and mammals, intrinsic apoptosis is considered to be ancestral and conserved among animals, and depends on shared initiators; caspase-9, Apaf-1 and Bcl-2. However, the biochemical role of mitochondria, the pivotal function of cytochrome c and the modality of caspase activation remain highly heterogeneous and hide profound molecular divergences among apoptotic pathways in animals. Uncovering the phylogenetic history of apoptotic actors, especially caspases, is crucial to shed light on intrinsic apoptosis evolutionary history. Here, we demonstrate by phylogenetic analyses, that caspase-9, the fundamental key of intrinsic apoptosis, is deuterostome-specific, while caspase-2 is ancestral to bilaterians. Our analysis of Bcl-2 and Apaf-1 confirm heterogeneity in functional organisation of apoptotic pathways in animals. Our results support emergence of distinct intrinsic apoptotic pathways during metazoan evolution.

## INTRODUCTION

Apoptosis, regulated cell death defined by a set of morphological features and dependent on caspases, occurs during metazoan development, tissue homeostasis, and regeneration^1–3^. Pioneering works in *Caenorhabditis elegans* established a molecular network of apoptosis decision, execution, engulfment-degradation, and more fundamentally the first description of intrinsic apoptosis, previously known as the mitochondrial apoptotic pathway^4–6^. Next, investigation of apoptotic cascade key components from *Drosophila melanogaster* and mammals imposed the paradigm that the intrinsic apoptotic “molecular program” is conserved throughout animal evolution^4,5,7–9^. Intrinsic apoptosis depends on the key initiator caspase-9 (named Ced-3 and Dronc in *Caenorhabditis* and *Drosophila*, respectively), the activator Apaf-1 (named Ced-4 and Dark in *Caenorhabditis* and *Drosophila*, respectively) and the Bcl-2 multigenic family^4,8^. However, recent research has revealed evolutionary divergences between these models, with major functional diversifications^10,11^. Importantly and independently of functional approaches, the phylogenetic history of the signal transduction actors, mostly the caspase family, is the Rosetta Stone to shed light on intrinsic apoptosis evolution. Here, we investigate the evolutionary genetic history of the major actors of intrinsic apoptosis. We conducted phylogenetic analyses of initiator CARD-caspases, Bcl-2 multigenic family, and Apaf-1 proteins from major animal phyla and describe their evolutionary patterns. Despite common functional similarities, these actors, especially initiator caspases, are engaged in relatively independent evolutionary histories that involve different signalling pathways. Consequently, we consider the structural similarities of intrinsic apoptotic components to be the result of convergent evolution among animals. Convergent evolution is defined here as a set of patterns (i.e. named intrinsic apoptosis) produced by similar but not identical processes (i.e. pathways driven by at least two groups of paralogous genes). Thus, the classical animal models (nematode, fly, mouse) which were used to establish a unified concept of intrinsic apoptosis are markedly distinguished by distinct molecular origin of their apoptotic actors but also by major functional divergences.

## RESULTS AND DISCUSSION

### Initiator caspases of intrinsic apoptotic pathways are not homologous genes

Initiation and execution of apoptotic signalling pathways are fundamentally linked to the complex diversification of caspases which are widely distributed among all metazoan phyla^12–15^.

Caspases are a class of proteases composed of three protein domains; the pro-domain, the small P10, and the large P20^16,17^. Initiator caspases, specific to each apoptotic pathway, have a Caspase Recruiting Domain (CARD) pro-domain or two Death Effector Domains (DED) pro-domains. Intrinsic and extrinsic apoptosis are distinct and involve specific initiators which are caspase-9 (CARD pro-domain) or caspase-8 or −10 (DED pro-domains), respectively. Both pathways trigger activation of common executioner caspases, leading to apoptosis execution^4,18^.

Due to the pivotal role of caspase-9 in initiating intrinsic apoptosis, our main analyses focused on caspases having a CARD pro-domain (CARD-caspases). We confirmed their distribution in most animal phyla and reconstructed their phylogenetic relationships (Figure 1, Table S1). Regardless of the phylogenetic methodology used for the analysis, two strongly supported (PP> 0.99) monophyletic groups were identified: caspase-9 and caspase-2 (Figure 1, Figure S1). A third and more heterogeneous group named here [Inflammatory Caspases + Caspase-Y] is monophyletic in Bayesian analysis, the method which produced the most likely topology (Table S2), and splits into two groups in maximum likelihood estimation (Figure 1, Figure S1). Although relationships among bilaterian clades differ depending on the methodology employed, monophyly of caspase-9 and caspase-2 families remains robust. They group together with Inflammatory Caspases and Caspase-Y in a clade strictly diversified within bilaterian animals (PP = 0.87; BS>90). The sequences of non-bilaterians [Cnidaria + Ctenophora + Placozoa] form a divergent paraphyletic group named here Caspase-X (Figure 1). Switching between metazoan putative “basal” out groups [i.e. *Amphimedon queenslandica* (Porifera) or *Pleurobrachia bachei* (Ctenophora)] does not affect caspase topology (Figure S2). All bilaterians sequences, caspase-9, caspase-2 and [Inflammatory Caspases + Caspase-Y] remain monophyletic (Figure S2).

**Figure 1:**
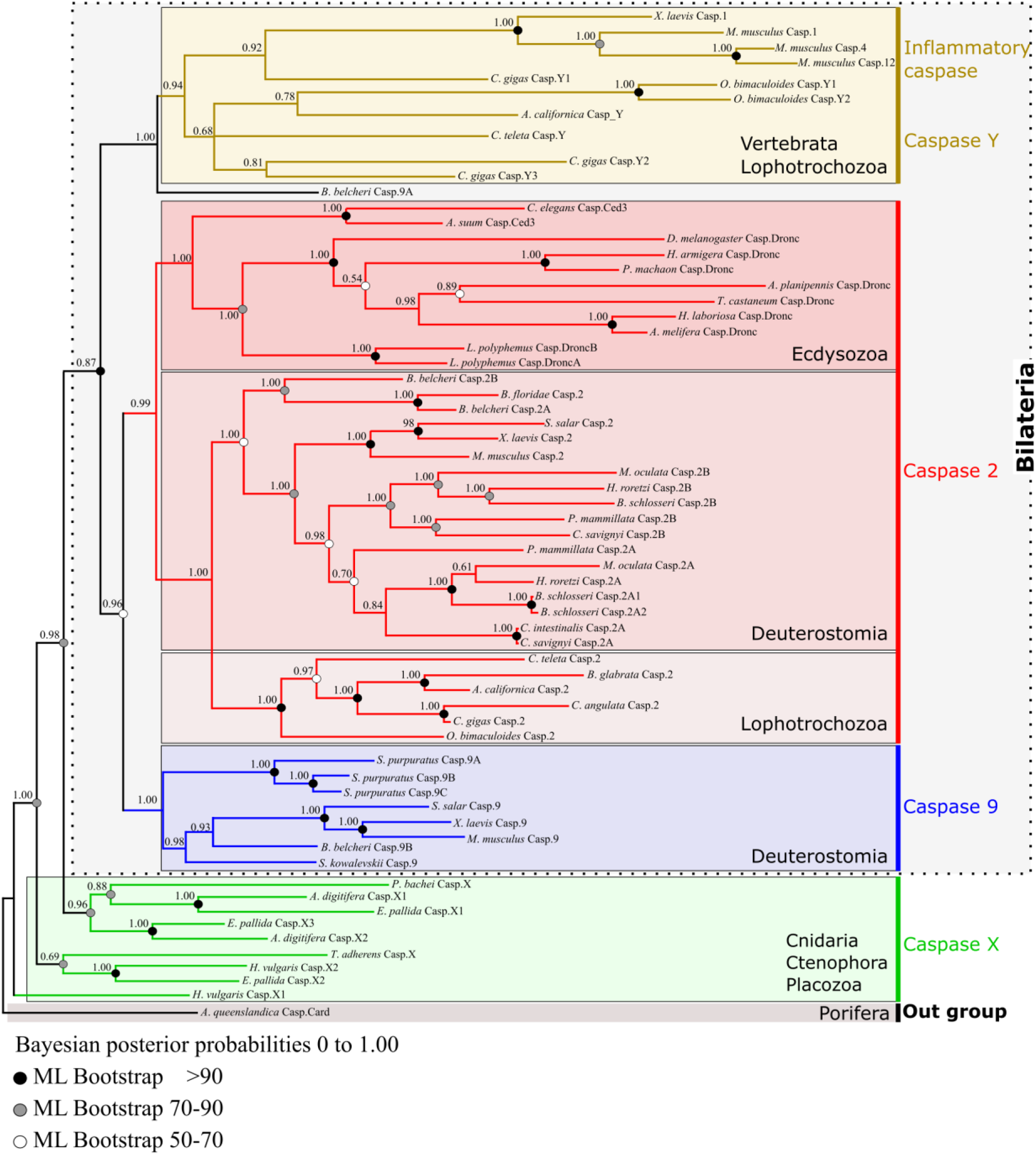
Topology of metazoan CARD-caspase phylogeny obtained by Bayesian inference using full sequences alignment. Three strongly supported monophyletic groups are identified: caspase-9, caspase-2 and a more heterogeneous group named here [Inflammatory Caspases + Caspase-Y]. Together they form a clade strictly corresponding to bilaterian animals. Caspase-2 is widely distributed among bilaterians [Deutostomia + Ecdysozoa + Lophotrochozoa/Spiralia] reflecting an ancestral origin. Conversely, caspase-9 is strictly restricted to deuterostomian animals. Sequences of non-bilaterians (Cnidaria + Ctenophora + Placozoa) form divergent paraphyletic groups. The selected outgroup is the unique CARD-caspase of Porifera *Amphimedon queenslandica*.

Caspase-2 appears to be widely distributed among bilaterians and sequences evolution is congruent with the bilaterian groups [Deuterostomia + Ecdysozoa + Lophotrochozoa/Spiralia] reflecting its ancestral origin. Surprisingly caspase-9 is strictly restricted to deuterostome animals. This distribution can be interpreted either as the early emergence of caspase-9 in all bilaterians followed by a secondary loss in protostomes, or as a specific derived character of deuterostomes. The diversity of the genomes explored, and their depth of sequencing unambiguously confirms this result. More specifically, the robustness of the deuterostome caspase-9 clade was explored (Figure S3), and its specific diversification among [Vertebrata + Cephalochordata + Echinodermata + Hemichordata] was confirmed. However, the relative position of the cephalochordate *Branchiostoma belcheri* 9A paralogous gene remains unstable. Consistent with previous studies on the ascidian *Ciona intestinalis*^19^, the five representatives of ascidian genomes studied are devoid of any caspase-9, likely due to a unique secondary loss in urochordates, the sister-group of vertebrates^20^.

Unexpectedly, *Drosophila* Dronc and *Caenorhabditis* Ced-3 are distinctly identified as orthologous (i.e. coming from speciation) to caspase-2 of vertebrates, and not to caspase-9, as was previously reported and largely accepted^21–28^. All Dronc/Ced3 proteins of insects, horseshoe crab (Xiphosura), and nematodes form a strongly supported monophyletic clade (PP = 1) revealing single gene conservation without duplication (except horseshoe crab). Thus, the caspase-2 clade of ecdysozoans is the sister group of caspase-2 clade of [Lophotrochozoa + Deuterostomia]. The absence of identifiable caspase-2 in echinoderms and hemichordates (i.e. Ambulacraria) can probably be interpreted as a clade-specific derived loss.

Conversely, the clade [Caspase-Y + Inflammatory Caspase] appears to be restricted to [Vertebrata and Lophotrochozoa].

To test evolutionary history and shuffling of different domains, we inferred recombination rate from CARD-Caspase coding sequences (reflecting intra-codon recombination). We investigated multiple alignments of domains sequences using the approximate Bayesian computation (ABC) approach based on coalescent evolutionary history of protein sequences, i.e. the framework *ProteinEvolverABC*^29^. The remarkable low level of recombination rate (Rho mean = 2.0328712) we estimated during CARD-Caspase family evolution rule out potential phylogenetic reconstruction artefacts (Table S3). As illustrative examples, this low recombination rate is comparable to those observed for some Coronaviruses protein families such as NS12.7 or astroviruses viroporins probably related to the maintenance of viral pathogenicity^29–31^. The multiple and essential roles played by the CARD-Caspase family within the cell is likely one factor explaining the extremely low recombination rate that can be observed.

Taken globally, our analysis shows that the traditionally considered conserved initiator caspases of intrinsic apoptosis are not orthologous, but are divergent genes putatively issued from ancestral duplication (i.e. paralogous genes) among bilaterians.

### The caspase activation regulator apoptosome is structurally divergent in metazoans

In intrinsic apoptosis, initiator caspase activation depends on their recruitment by a pivotal, and considered shared component, the apoptosome platform^4,10^. Apoptosome formation results on critical protein binding between *Caenorhabditis* Ced-4, *Drosophila* Dark, and human Apaf-1 with their respective initiator caspases Ced-3, Dronc, and procaspase-9, respectively^10^. CARD and other domains (NOD, arm) are highly conserved in Apaf-1, Dark and Ced-4 proteins^32–34^. However, excepted the majority of nematode species^11^, Apaf-1 possesses WD40 repeats at its C-terminus which bind to cytochrome c. This binding is required in mammals for Apaf-1 oligomerisation and apoptosome formation^34,35^, while Ced-4 and Dark do not require cytochrome c for their general assembly into an apoptosome^10,36^. Taken with major structural and regulating assembly differences between the octameric Dark, tetrameric Ced-4 and heptameric Apaf-1 apoptosomes, it reveals evolutionary divergence between animal apoptosomes formation and procaspases activation mechanisms, probably differentially modulating the cell death execution pathway^37^.

We conducted exhaustive studies by reciprocal BLAST and phylogenetic analyses of Apaf-1 homologs and confirmed their conservation in the majority of metazoan phyla (Figure S4, Table S4)^32,33^. We conducted our analysis with all sequences issued from a previous study^33^ supplemented with ones we have specifically identified. However, we have differentiated in our study Apaf-1 genes sharing a CARD domain, and likely involved in apoptosis regulation, from Apaf-1-like devoid of CARD domains but which possess a panel of unusual domains. NB-ARC being the most conserved and shared domain by almost all Apaf-1 and Apaf-1-like genes among metazoans, we conducted the analysis on the NB-ARC alignment as it was previously done^33^.

Our topology presents two main clades that are common between Bayesian inference method and maximum likelihood estimation, both of which are inconsistent with species phylogeny. One of them grouped ecdysozoan sequences including Ced-4 and Dark in addition to several Apaf-1-like from the cnidarian *Nematostella vectensis* and the cephalochordate *Branchiostoma floridae*, while the other one contains deuterostomian and non-bilaterian sequences. Unexpectedly, deuterostomians are not monophyletic but split into two groups, suggesting classic Long Branch attraction artifact and a very fast sequence evolution. Remarkably, Apaf-1 was absent from all available urochordate and mollusc genomes, in accordance with previous work^19,33,38–41^, and from *Capitella teleta* (Annelida). Even if differences in the quality of the genomes might account for some of the putative losses, these results suggest independent losses in urochordates and lophotrochozoans or, a less parsimonious hypothesis, complex and independent modular protein evolution from the common ancestor of metazoans (Figure 2). In non-bilaterians, the Apaf-1 gene family has not been found in the *Pleurobrachia pileus* genome (Ctenophora), but seems present in other early diverging metazoan phyla [Porifera, Cnidaria and Placozoa] (^33,42,43^, our data).

**Figure 2.**
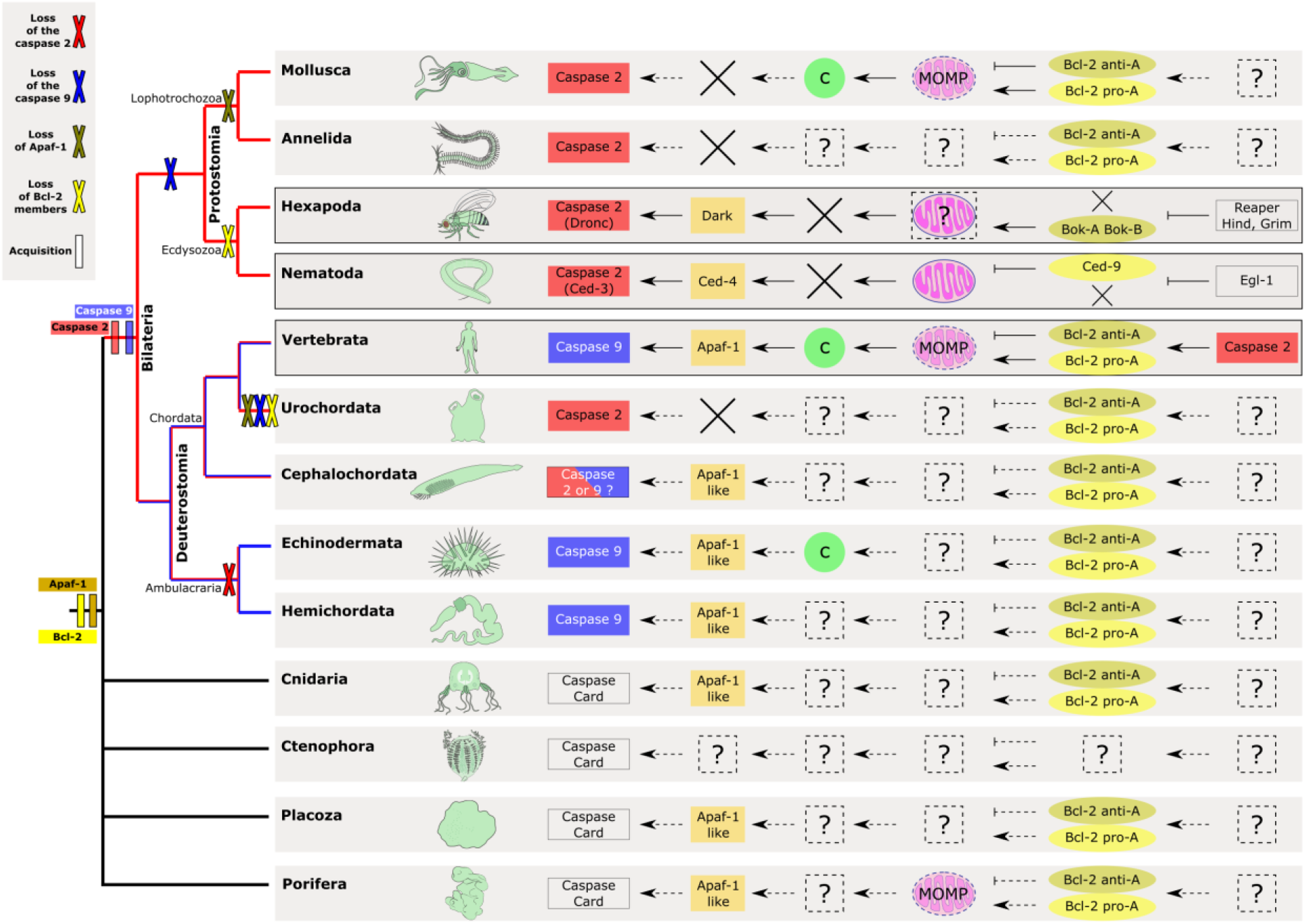
Reconstruction of convergent hypothetical intrinsic apoptotic pathways among metazoans according to molecular actors detected and identified in their genomes. Variability of intrinsic apoptotic pathways among animals emerged from convergences and recruitment of distinct actors with independent evolutionary history. Caspase-2 is bilaterian-specific and the initiator of ecdysozoans intrinsic apoptosis. Caspase-9 is restricted to deuterostomes and the specific initiator of mammalian intrinsic apoptosis. Deuterostomes exhibit several losses (i.e. caspase 2 in cephalochordates, caspase 9 in urochordates) or duplication (i.e. caspase 9 in echinoderms), highlighting a putative evolutionary flexibility in apoptotic pathway establishment. Mitochondrial functions diverge among phyla and cytochrome c release thanks to mitochondrial outer membrane permeabilisation (MOMP) is specific only to mammals and possibly echinoderms. The convergent evolutionary histories reflect a probable phylum specific adaptive process leading to parallel evolution of mitochondrial apoptotic pathways observed among animals.

Importantly, a previous analysis from several genomes (cnidarian, nematode, fly, amphioxus, sea urchin, human) clearly identified three well-defined independent clades of genes among metazoans, highlighting that Ced-4, Dark and Apaf-1 are not homologous genes between ecdysozoans and vertebrates^33^.

Despite its fundamental role in the apoptotic cell death pathways and also in non-apoptotic functions^44–46^, diverse metazoan phyla have independently experienced adapter protein Apaf-1 losses. Apaf-1 with CARD-domain, likely involve in apoptosis, are under strong purifying selection as illustrated by the metric non-synonymous/synonymous substitution rate ratio (dN/dS) close to 0 (model M0; ωθ = 0.07), perhaps promoted by their crucial and specific role as caspase activator. Consequently, we hypothesise a relaxation of functional constrains on unusual Apaf-1 molecules (i.e. deprived of Card-domain, and probably not involved in oligomerisation processes) that specifically drive the diversification of Apaf-1/Apaf-1-like paralogous genes in the cnidarian *Nematostella vectensis* and the cephalochordate *Branchiostoma floridae*^33^.

These elements argue in favour of distinct evolutionary trajectory between species and convergence of the modality of apoptosome formation, which could explain functional analyses showing that its formation and the subsequent caspase activation differs among taxa^11^.

### The regulation of apoptosis by the Bcl-2 family is divergent among metazoans

Intrinsic apoptosis is ultimately regulated by the Bcl-2 proteins, composed of several Bcl-2 homologous (BH) domains^47,48^. In mammals, the balance between pro-survival (four BH1-BH4 domains) and pro-apoptotic proteins (Bax/Bak/Bok and BH3-only) of Bcl-2 controls initiation of intrinsic apoptosis. Conformational changes of the three-dimensional structures and interactions between Bcl-2 actors enables the assembly of pore-like structures controlling MOMP^49^.

Multiple sequence alignments of metazoan Bcl-2 family proteins (Figure S5, Table S5) (but with over-represented chordates reflective of greater availability of vertebrate genomes) confirm the wide distribution and early origin of Bcl-2 in Metazoa^11,32,48,50^. Proteins clustered into five monophyletic groups, consistent with classical functional Bcl-2 classification (i.e. three ‘pro-apoptotic’ clades: Bok, Bak, Bax and two less supported pro-survival - ‘anti-apoptotic’ - groups: Bcl-2/W/XL and a more complex Bcl-B/Mcl-1/Bfl-1 clade) (Figure S5). However, relationships among the five well-supported groups were not clearly resolved despite the use of various methodologies and outgroup selections (Figure S5A-B). Consequently, each of the five Bcl-2 clades seems conserved across evolution, and homology (orthology) relationships inside each clade can be established. However, due to the impossibility of addressing the relationship between these five monophyletic groups (Bok, Bak, Bax, Bcl-2/W/XL, Bcl-B/Mcl-1/Bfl-1), it seems difficult to hypothesises their evolutionary pattern, history, and determine a potential ancestral group. In addition, very divergent sequences from molluscs such as *Biomphalaria glabrata* (Bcl-like2, Bcl-like3), from the cnidarian *Hydra vulgaris* (Bcl-like1) or from the urochordate *Ciona intestinalis* (Bcl-like1) were not assigned to a particular class.

*Caenorhabditis* is deprived of pro-apoptotic Bcl-2 and possesses only the pro-survival Ced-9 (homolog to vertebrates Bcl-2/w/xl), and two BH-3 only proteins (Egl-1 and Ced-13). Conversely, only two pro-apoptotic Bok-like paralogs (Debcl and Buffy) were present in *Drosophila* (Figure 2, Figure S5), but their functions remain unclear^51^. Presence of both anti- and pro-apoptotic Bcl-2 in molluscs underlines the divergent particularities observed within protostomes. Contrarily, similarities in Bcl-2 family composition are observed within some deuterostomes (mammals and echinoderms).

Finally, if multiple Bcl-2 genes were acquired early in metazoan evolution, and despite conservation of almost homologous genes, key differences accumulate, making initiation mechanisms in intrinsic apoptotic signalling pathways divergent among animals.

### Apoptotic mitochondrial pathways are divergent among metazoans

Functional evidence shows that caspase-2 members play a critical role in various cell death processes, but are independently involved in a range of non-apoptotic functions, including cell cycle regulation, DNA repair and tumour suppression^52–55^. This implication of caspase-2 in a myriad of signalling pathways, and its ability to interact with a panel of molecules, demonstrates its functional versatility^52,53,56–59^. As previously reported, our phylogenetic analyses corroborate the wide distribution of caspase-2 in bilaterians and suggest a probable ancestral multifunctionality.

The major structural and functional similarities that led to an erroneous interpretation of the phylogenetic position of Ced-3 and Dronc underlies a probable common evolutionary origin of caspase-2 and −9 genes. We hypothesised here that caspase-9 originates from bilaterians-specific duplication of a caspase-like ancestor gene, followed by a loss of caspase-9 in protostomians (Figure 2). Among deuterostomians the two families of paralogs have been preserved in vertebrates and probably in cephalochordates (Figure 1, 2). Caspase-2 retains multifunctional activity, and in mammals can interact with the PIDDosome platform containing P53, adapter molecule RAIDD, and signalling complex DISC, activating both extrinsic and intrinsic apoptosis, and DNA damage pathways^58,60–62^. Conversely, the caspase-9 gene underwent a functional divergence in connection with its specialisation in allosteric interactions with the apoptosome^18^.

Due to its pivotal role as a mediator of genomic stability through involvement in cell proliferation, oxidative stress, aging and cell death, the molecular divergence of the caspase-2 gene is highly constrained during evolution, probably because destabilisation of any signalling cascade is sufficient to initiate tumorigenesis. Indeed, purifying selection is likely important during caspase-2 evolution as illustrated by a dN/dS close to 0 (model M0 ωθ = 0.07)^63,64^. However, during the radiation of deuterostomes, both presence of caspase-2 and - 9 certainly facilitated the loss of one of them (cf. DCC Force’s model - Duplication-Degeneration-Complementation model)^65–67^. Interestingly, a caspase-2 duplication co-occurred with the loss of caspase-9 in urochordates that may indicate a clade-specific relaxation of purifying selection on caspase-2 gene (Figure 1, Figure 2). Similarly, caspase-2 has been lost, or strongly diverged, while caspase-9 presents a *de novo* relative expansion in *Strongylocentrotus purpuratus* (Echinodermata) and *Saccoglossus kowalevskii* (Hemichordata). These expansions certainly result from a less powerful purifying selection of caspase-9, showing a higher dN/dS (model M0 ωθ = 0.22) than caspase-2.

Due to generalised gene losses in ecdysozoans and in comparison with other bilaterians, *Caenorhabditis* and *Drosophila* apoptotic pathways are generally considered simpler than those of vertebrates^17,68^. However, comparing to mammals, ecdysozoans have pathways organised around different paralogous genes in addition to a smaller number of genes present. The absence of orthologous genes relationships results in a very different structural organisation of apoptosome platforms but also generates important functional divergences (i.e. mechanisms of regulating assembly, CARD-CARD interactions with procaspases)^10^.

Consistent with mammalian caspase-2 functions, Ced-3 has both initiator as well as executioner activities^58,69–71^. Involvement of Dronc in various processes such as compensatory cell proliferation, inhibition of cell migration or spermatid differentiation, brings this protein closer to the functionalities of mammalian caspase-2^72,73^. Due to its interaction with Dark (Apaf-1 paralog), the only CARD-caspase in *Drosophila* (Dronc) has been erroneously classified as caspase-9^8,74^. Unlike the organisation of the mammal apoptosome, both *Caenorhabditis* and *Drosophila* present neither MOMP nor the release and necessity of cytochrome c to activate Ced-3 and Dronc via Ced-4 and Dark, respectively^5,68^.

Like other protostomes, molluscs are devoid of caspase-9 but caspase-2 has been identified in bivalves and was suspected to function in “a caspase-9-like manner”^75^. Hence, caspase-2 seems involved in apoptotic processes in molluscs, but more surprisingly, despite the absence of caspase-9 and Apaf-1 (and thus, of a mammalian-like apoptosome), this peculiar pathway is surprisingly associated with cytochrome c release^38,41,76–79^. The complexity of intrinsic apoptosis in molluscs seems to be important, but divergent from what was observed in ecdysozoans or vertebrates, with a putative expansion of caspases that participate both in immunity, stress responses, and apoptosis (Figure 2)^41,75,80–83^.

Unexpectedly, mammals present an almost unique case (possibly with the cephalochordates) in which both caspase-2 and caspase-9 are conserved and involved in apoptosis. This putative functional redundancy (i.e. recruitment, autoactivation or transactivation, homodimerisation and subsequent interchain proteolytic cleavage) undoubtedly led to the functional specialisation observed for caspase-9. This may results from the specificity of the mammalian mitochondrial pathway and the non-apoptotic function of caspase-9^84–87^. Finally, echinoderms seem to uniquely have an intrinsic apoptosis similar to mammals, with a caspase-9, Bcl-2, Apaf-1, and a MOMP with cytochrome c release (Figure 2)^7,88^.

Although we can envisage a weak parsimonious scenario that showcases a common ancestral apoptotic pathway in deuterostomes (but implying independent secondary losses in hemichordates, cephalocordates, and urochordates), the similarities observed between echinoderms and mammals more probably reflect “functional convergences” based on independent recruitment of apoptotic actors.

## CONCLUSION

The apoptotic networks of *Caenorhabditis* and *Drosophila* do not exemplify ancestral conditions from which mammalian-grade apoptotic complexity emerged but are on the contrary, and as suggested recently, the result of a derived condition specific to ecdysozoans among animals ^32,80,89,90^.

The core components of intrinsic apoptotic pathways, especially initiator caspases and the apoptosome platform, are not ancestral in metazoans. Our phylogenetic analyses highlight an unexpected evolutionary history: while the bilaterian caspase-2-mediated apoptotic toolkit emerged ancestrally and remains multifunctional, the caspase-9 mediator of the mammalian apoptosome is specific to deuterostomes.

The major functional divergences in mitochondrial apoptotic pathways observed in animals^70,91–93^ mainly originated in the recruitment of paralogous actors from the same multigenic families. The evolution of the different pathways underlined by non-homologous molecular actors could be the reflection of putative adaptive processes or constrains specific to each taxon, which ultimately led to divergent evolutionary histories. Interestingly, the richness of the apoptotic genetic repertoire was suggested to be linked to the persistence of stem cells in adults from different phyla^94,95^.

Finally, mitochondria-mediated apoptosis, like other programmed cell deaths, has likely evolved before and throughout metazoan diversification to shape developmental processes, immune response, or to adapt cellular environment to environmental constraint.

## MATERIAL AND METHODS

### Sequence dataset construction

Putative metazoan caspases with a CARD pro-domain (CARD-caspases) were identified using tBLASTn and BLASTp searches with human caspases, Ced-3, and Dronc as queries on NCBI, ANISEED (ascidians), EchinoBase (*Strongylocentrotus purpuratus*), and neurobase.rc.ufl.edu (*Pleurobrachia bachei*) databases, and followed by reciprocal BLAST. After identification of CARD-caspases in target species, sequences were added as queries to conduct BLAST searches in close relatives (i.e. identified CARD-caspases of *Crassostrea gigas* were used as queries to search in other molluscs). Genomes and transcriptomes of species’ were downloaded when possible, to search a second time. Sequences with an e-value inferior to 1e-10 were retained. All identified sequences were analysed with ScanProsite (ExPaSy)^96^ and InterProScan (EMBL-EBI)^97^ to verify the presence of specific caspase domains. Another sequence was added to verify its identification proposed as a caspase-2 in the literature: *Crassostrea angulata* caspase-2^79^. Caspase family proteins are short (containing the large common P20 and the small P10 domains) with a high number of genes per species which rapidly limits the relevance of the phylogenetic analyses. To reduce artefact branching and unreadable topologies, and to maximize phylogenetic diversity across Metazoa, the dataset was built using CARD-caspase gene repertoires of selected species. A full list of all caspase sequences is provided in Table S1. Taxon and genes were selected to have a wide diversity of major metazoan phyla and to have an equilibrium inside and between each group. We choose representative species for each phylum and took care to don’t unreasonably increase the number of sequences by redundant choice. We privileged species where we detected caspases with CARD domains, DED domains, and executioner caspases, suggesting a high quality of sequences allowing identification of all members of the caspase family. For each taxon, all caspases having a CARD pro-domains have been include in the analysis.

Identification of metazoan Apaf-1 was made using tBLASTn and BLASTp using human APAF-1, nematode Ced-4, and fly Dark as queries on NCBI, ANISEED (ascidians), and neurobase.rc.ufl.edu (*Pleurobrachia bachei*) databases, and followed by reciprocal BLAST. Potential resulting sequences were analysed with ScanProsite and also InterProScan. A full list of all Apaf-1 sequences is provided in Table S4. Apaf-1 data set comprise sequences from a previous study^33^ in addition to our new sequences.

Metazoan Bcl-2 were identified by using tBLASTn and BLASTp searches with human Bcl-2 as the query sequence on NCBI, and then again on downloaded genomes and transcriptomes, followed by reciprocal BLAST. All identified sequences were analysed with ScanProsite (ExPaSy) and InterProScan (EMBL-EBI) to verify the presence of BH domains. Due to short sequence length, BH3-only were not taken into account. A full list of all Bcl-2 sequences is provided in Table S5.

Multiple alignments of protein sequences were generated using the MAFFT software version 7^98^ with default parameters and also Clustal Omega^99^ to verify the congruence of the different alignments. All sequences were then manually checked in BioEdit 7.2 software^100^ to verify the presence of specific domains previously identified. Gblocks version 0.91b^101^ was used to remove vacancies and blur sites. Final alignments comprise of 230, 235, 147, and 306 amino acids for metazoan CARD-caspases alignment, deuterostomian CARD-caspases alignment, metazoan Bcl-2 alignment, and Apaf-1 alignment, respectively. Apaf-1 alignment was done on the NB-ARC domain only.

### Phylogenetic analysis

Phylogenetic analyses were carried out from the amino-acid alignment with Maximum-Likelihood (ML) method in PhyML 3.1^102^, a combined ML tree search with 1000 bootstrap replicates was produced and then visualised using Seaview^103^. The best amino-acid evolution models to conduct analyses were determined using MEGA11^104^ and determined to be WAG for sequences of CARD-caspases alignments (CARD, P20, P10) and LG for both Bcl-2 and Apaf-1 alignments.

Bayesian analyses were performed using MrBayes (v3.2.6)^105^ under mixed model. For each analysis, one fourth of the topologies were discarded as burn-in values, while the remaining ones were used to calculate posterior probability. The run for metazoan CARD-caspases alignment were carried out for 2,000,000 generations with 15 randomly started simultaneous Markov chains (1 cold chain, 14 heated chains) and sampled every 100 generations. The run for deuterostomian CARD-caspases alignment was carried out for 500,000 generations with 5 randomly started simultaneous Markov chains (1 cold chain, 4 heated chains) and sampled every 100 generations. The run for metazoan Apaf-1 alignment was carried out for 500,000 generations with 5 randomly started simultaneous Markov chains (1 cold chain, 4 heated chains) and sampled every 100 generations. The run for metazoan Bcl-2 alignment was carried out for 5,000,000 generations with 20 randomly started simultaneous Markov chains (1 cold chain, 19 heated chains) and sampled every 100 generations. ML bootstrap values higher than 50% and Bayesian posterior probabilities are indicated on the Bayesian tree.

The outgroup for the metazoan CARD-caspase phylogeny is the only caspase with a CARD pro-domain of the Porifera *Amphimedon queenslandica* (XP_003383519) (Figure 1, S4, S5) or the only one of the ctenophore *Pleurobrachia bachei* (Sb 2658116) (Figure S2). Analyses of CARD-caspases were made independently at the deuterostome scale with four different outgroups to test their effect on the stability of the topology: *i*) CARD-caspase-Y of the annelid *Capitella teleta* (ELT97848.1), *ii*) CARD-caspase-2 of the mollusk *Aplysia californica* (XP_005113266), *iii*) CARD-caspase-X2 of cnidarian *Hydra vulgaris* (NP_001274285.1) *iv*) CARD-caspase Ced-3 of the ecdysozoan *Caenorhabditis elegans* (AAG42045.1) (Figure S3). For the metazoan Apaf-1 phylogenies, the selected outgroup is Apaf-1 (XP_019855714.1) of Porifera *Amphimedon queenslandica* (Figure S4). For the metazoan Bcl-2 phylogenies, outgroups used to test their effect on stability of the topology are: *i*) Bcl-2-like1 (XP_003383425.1) and Bcl-2-like2 (XP_003387574.1) of Porifera *Amphimedon queenslandica* (Figure S5).

Topology tests between metazoan CARD-caspase phylogenies from Bayesian inference method and maximum likelihood estimation were evaluated using IQ-TREE^106^. We conducted Kishino-Hasegawa test^107^, Shimodaira-Hasegawa test, and approximately unbiased test^108^.

### Estimation of dN/dS and recombination rate

Codon aligned nucleotide sequences were built using PAL2NAL^109^ from clustalo aligned amino acid sequences and gene ORFs. The dN/dS ratio was calculated using EasyCodeML under settings “site model”, nuc-2, and standard icode^64^, from the codon aligned nucleotide sequences and newick format trees.

Co-estimation of recombination rate was conducted using ProteinEvolverABC^29^. We run the analysis on the same multiple protein alignment we used for the metazoan CARD-caspases phylogenetic analysis, including CARD, P20, and P10 domains. Analysis was conducted with 10 000 simulations and an acceptance rate of 0.2% to optimise the accuracy of the estimations (Beaumont 2010), and under WAG amino acid substitution model. Prior parameters were estimated by ProteinEvolverABC prior to the analysis such as Rho = Uniform (0, 3.0e^−6^) and Theta = Uniform (9.1e^−5^, 4.0e^−4^). Proportion of invariable sites was 0.026 and amino acid frequencies were fix.

## Acknowledgements

Authors particularly acknowledge Helen R Horkan (University of Galway, Ireland) for help in bioinformatics analysis and helpful comments, and Miguel Arenas (University of Vigo, Spain) for discussion and assistance with ProteinEvolverABC. Authors thank Sébastien Darras (Sorbonne Université, Banyuls-sur-Mer), Christine Vesque (Sorbonne Université, Paris), Jérôme Gros (Institut Pasteur, Paris), Sabine Hennequin (Sorbonne Université, Paris) and Uri Frank (University of Galway, Ireland) for helpful comments. GK was supported by a Ph.D. fellowship from the French Ministry of Education, Research and Innovation.

## Author Contributions

JP and EQ managed the project. GK made BLAST, phylogenetic and sequences analysis, and figures. GK and EQ wrote the manuscript.

## Data Availability

All data needed to evaluate the conclusions in this study are present in the paper and the Supplementary Materials. Any requests can be addressed to the corresponding author.

---

**Supplementary Figure 1:**
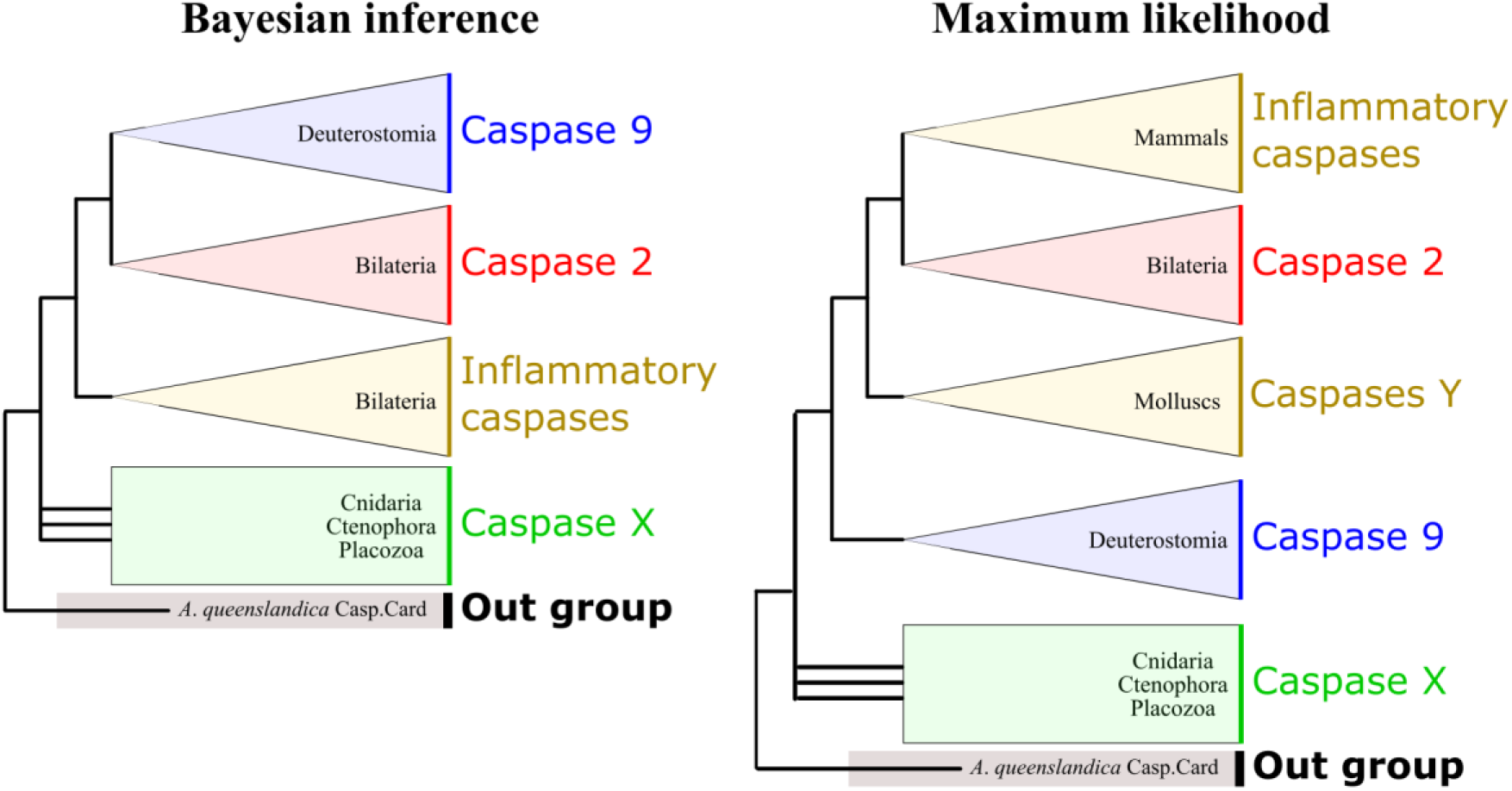
Monophyly of each CARD-caspase group is conserved using Bayesian inference and maximum likelihood analyses. Caspase-9, caspase-2 and [Inflammatory Caspases + Caspase-Y] groups remain conserved, confirming the bilaterian-specificity of caspase-2 and deuterostomian-specificity of caspase-9. Relationships among these three clades differ depending on the methodology employed. Divergent sequences of non-bilaterians animals (Cnidaria + Ctenophora + Placozoa) always form a paraphyletic group of caspases.

**Supplementary Figure 2:**
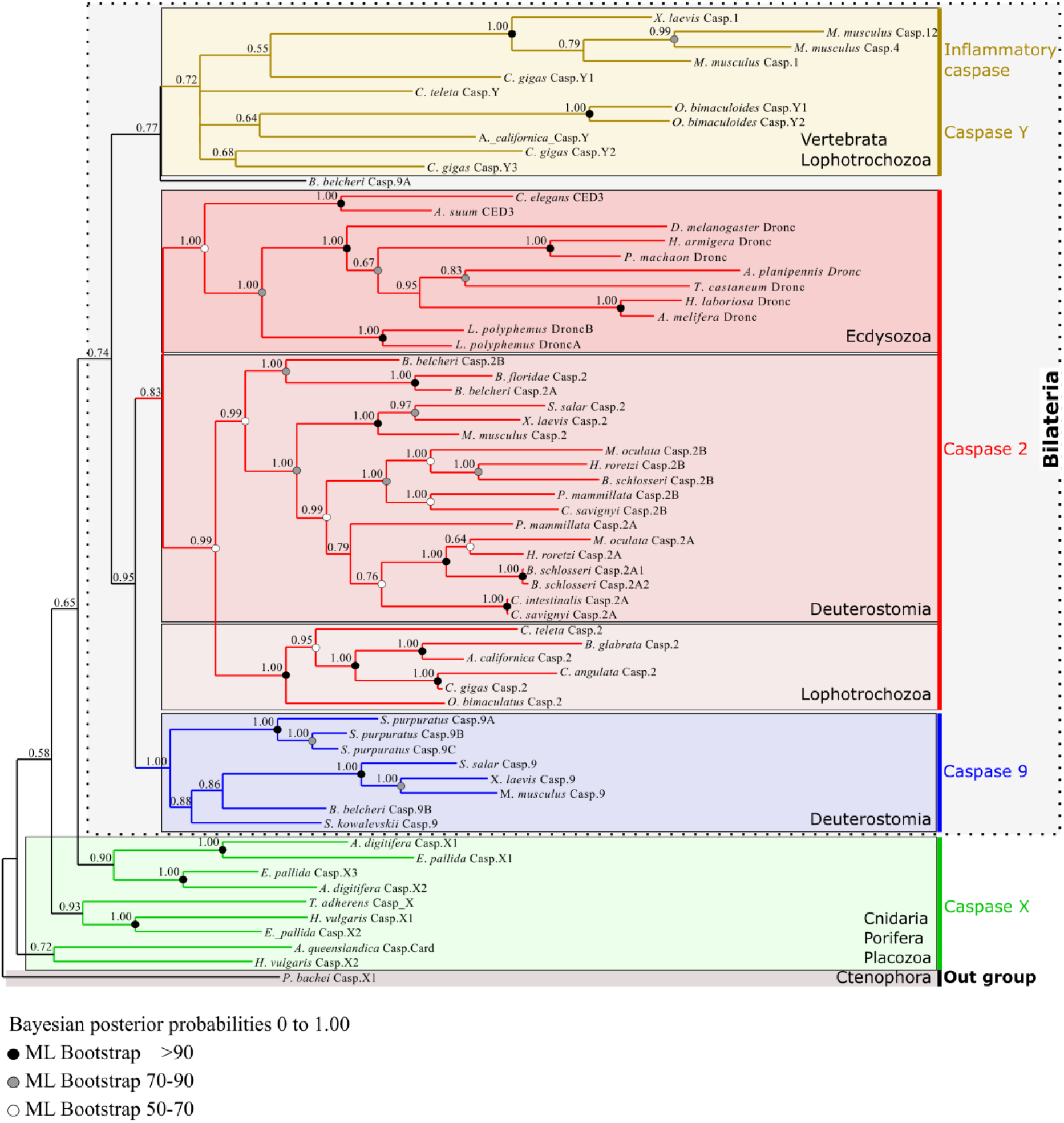
Topology of metazoan CARD-caspase phylogeny obtained by Bayesian inference. Selected outgroup is the unique CARD-caspase of Ctenophora *Pleurobrachia bachei*. The three strongly supported monophyletic groups identified, caspase-9, caspase-2 and [Inflammatory Caspases + Caspase-Y], are the same as when we used the unique CARD-caspase of the Porifera *Amphimedon queenslandica* as outgroup. Also, sequences from bilaterian animals remain monophyletic.

**Supplementary Figure 3:**
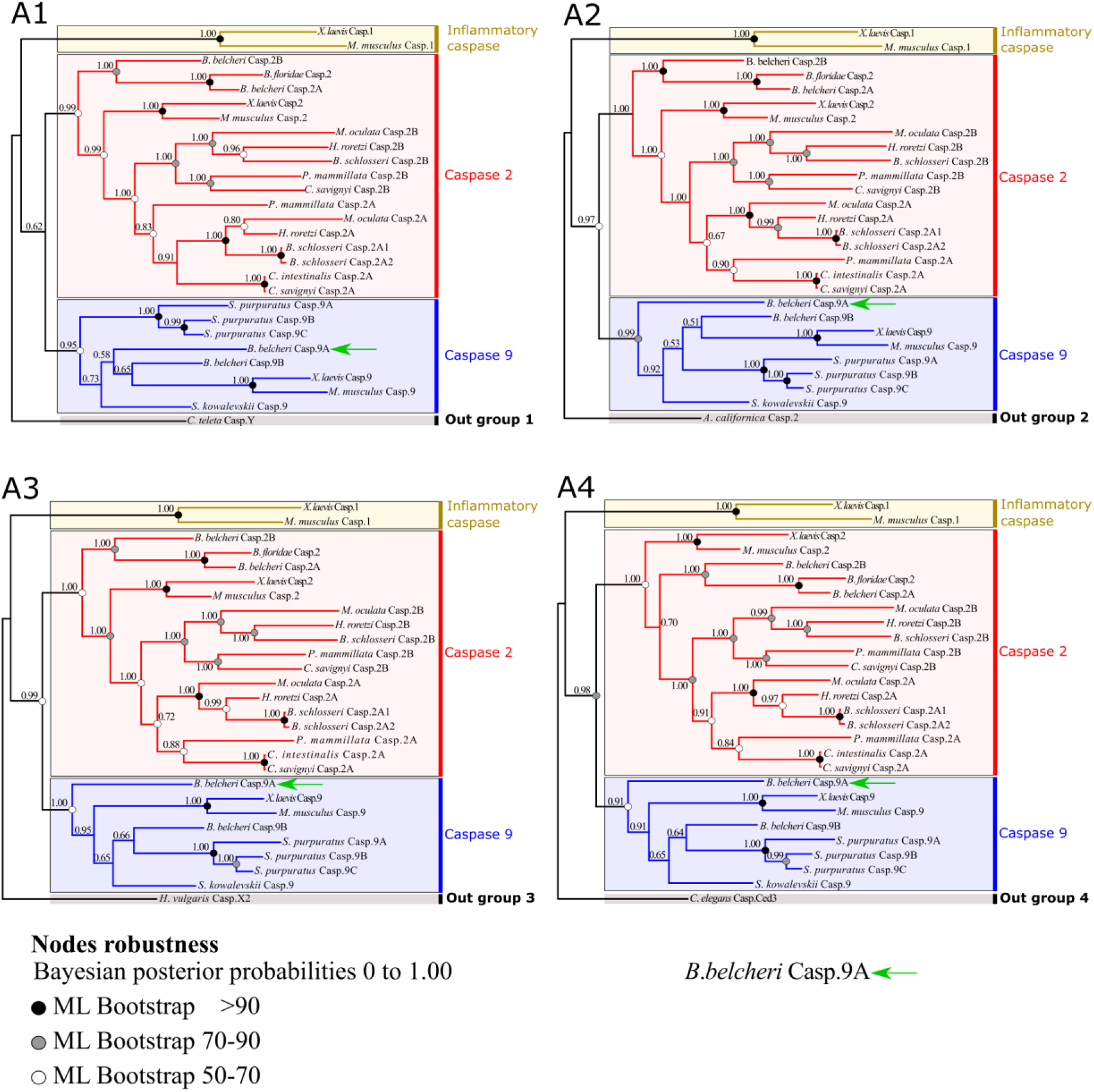
Phylogeny of CARD-caspases at the deuterostomian scale. Maximum likelihood and Bayesian inference methods produce similar topologies. Despite the unstable position of cephalochordate *Branchiostoma belcheri* caspase-9A (green arrow), it unequivocally appears to belong to the caspase-9 group. Numbers given correspond to posterior probabilities. We used respectively as outgroups *Capitella teleta* caspase-Y (ELT97848.1) (**A1**), *Aplysia californica* caspase-2 (XP_005113266) (**A2**), *Hydra vulgaris* caspase-X2 (NP_001274285.1) (**A3**), and *Caenorhabditis elegans* Ced3 (AAG42045.1) (**A4**).

**Supplementary Figure 4:**
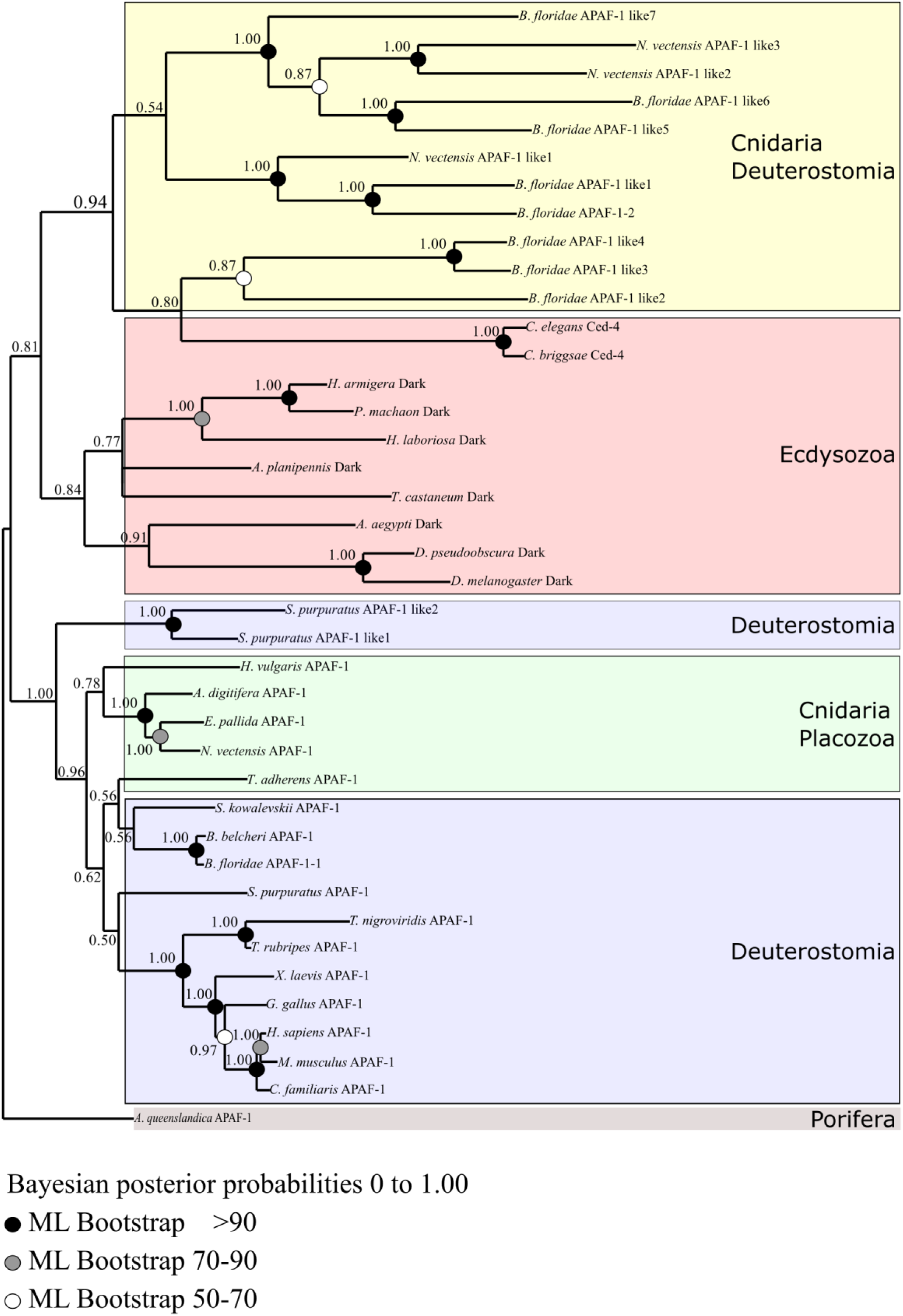
Phylogeny of Apaf-1 at the metazoan scale made with the NB-ARC domain alignment. Two monophyletic groups are identified: Apaf-1 of ecdysozoans, and apaf-1 of [Deutostomia + Cnidaria + Placozoa]. Globally, the topology is not consistent with the species phylogeny, and nodes are poorly supported. Among metazoans, Apaf-1 is characterised by high sequence divergence and fast independent evolution, making homology hypotheses difficult to establish. The selected outgroup is Apaf-1 of Porifera *Amphimedon queenslandica*.

**Supplementary Figure 5:**
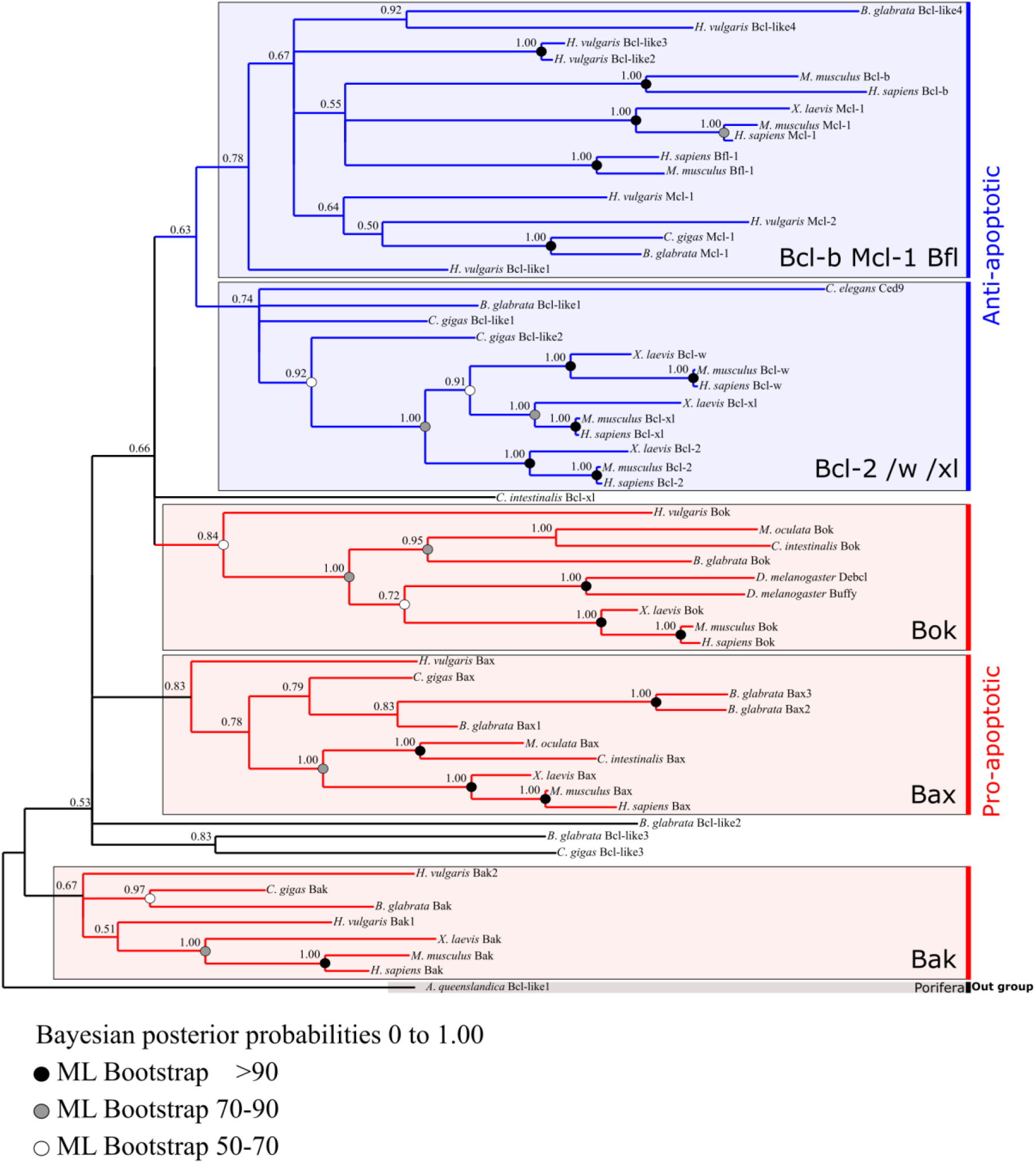
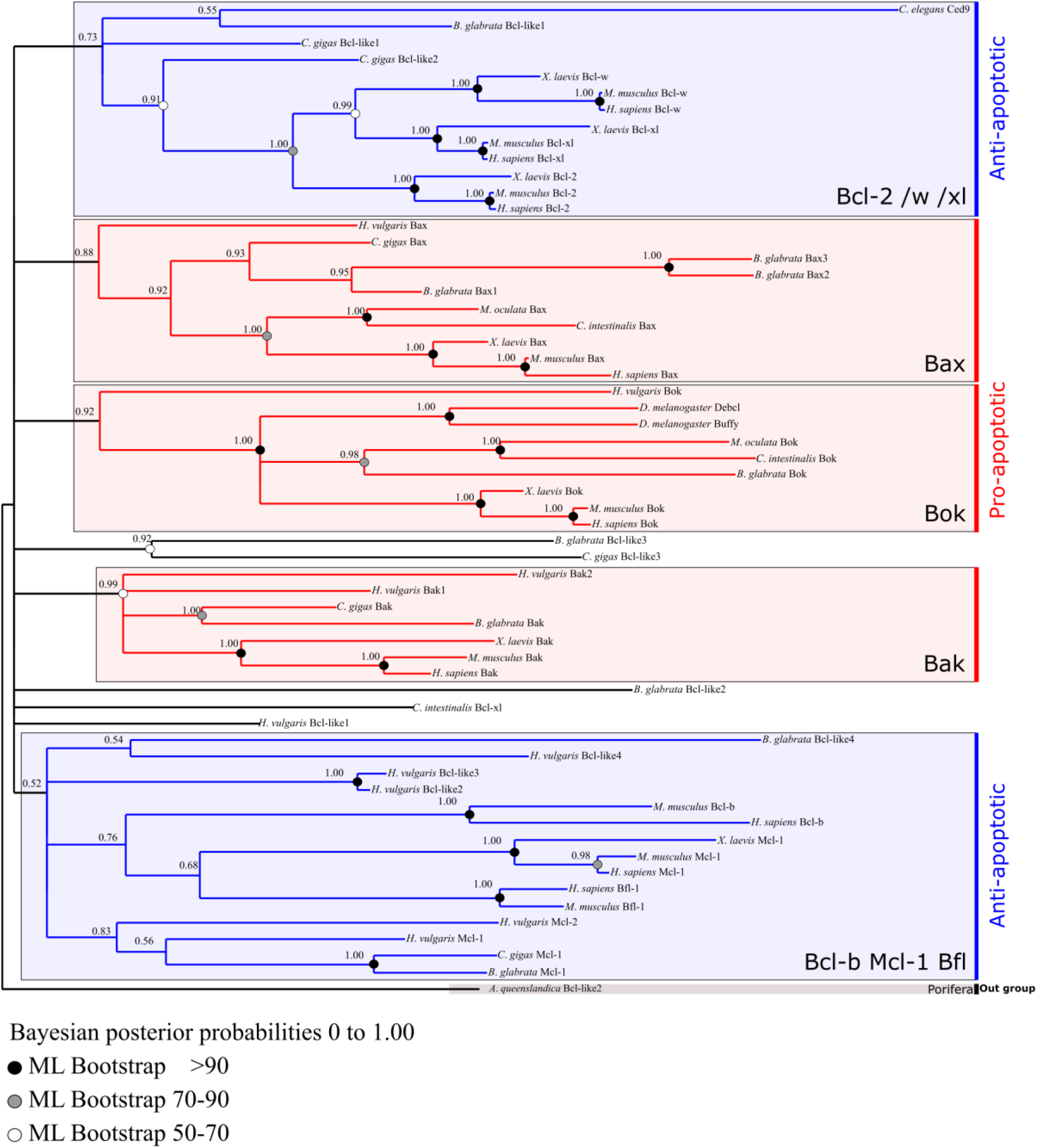
Topology of Bcl-2 family phylogeny from Bayesian inference and maximum likelihood at the metazoan scale using outgroups *Amphimedon queenslandica* Bcl-like 1 (XP_003383425.1) (**A**) and Bcl-like 2 (XP_003387574.1) (**B**). Bcl-2 proteins strictly clustered into five monophyletic groups: three ‘pro-apoptotic’ clades (Bok, Bak, Bax) and two less supported ‘anti-apoptotic’ groups (Bcl-2/W/XL) and a more complex (Bcl-B/Mcl-1/Bfl-1) clade. Each group includes bilaterian as well as non-bilaterian sequences which suggests a deep origin of this complex multigenic family. Clustering into these five groups is consistent between analyses, while relationships among them is not well resolved.

**Supplemental table 1:**
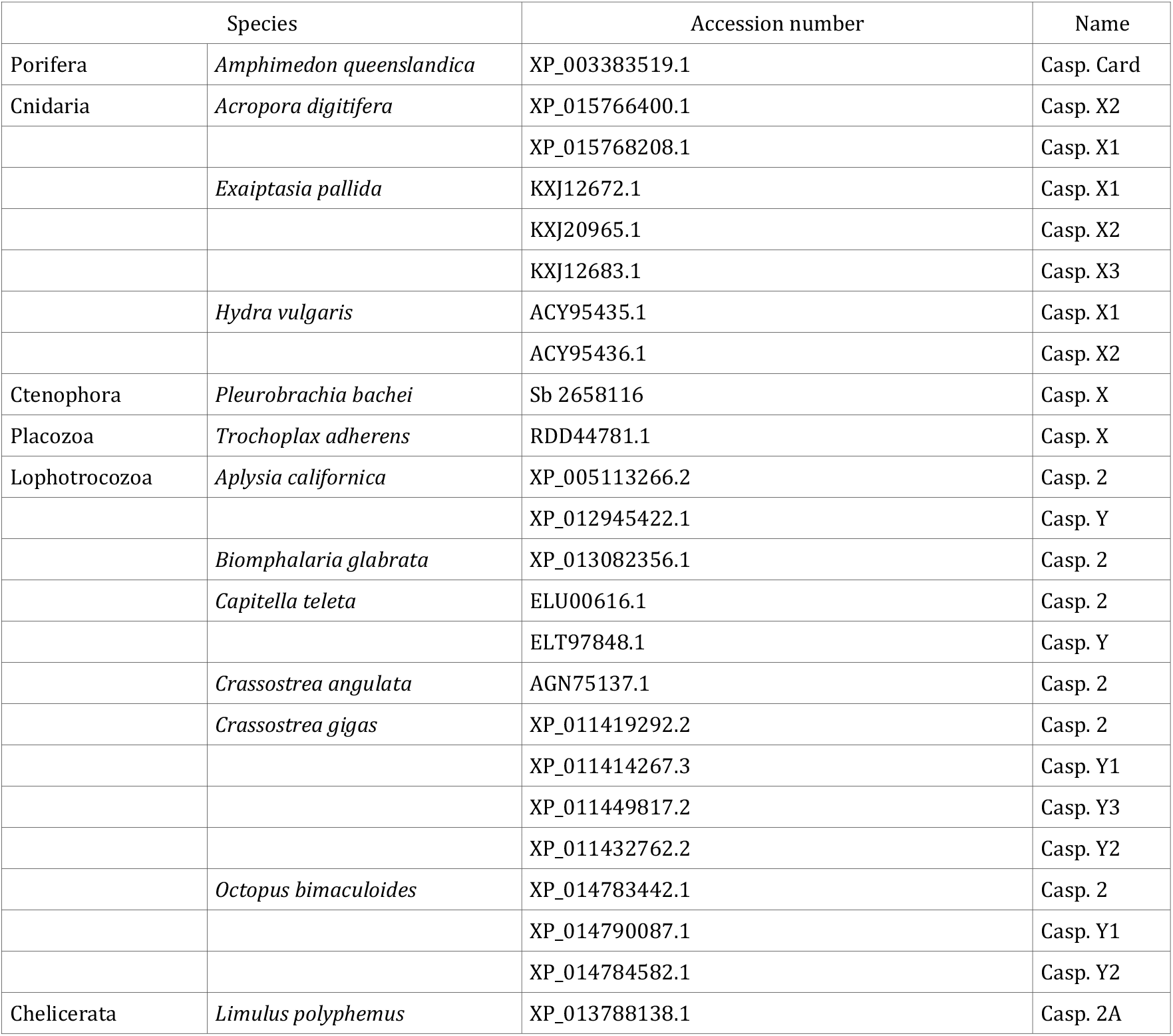
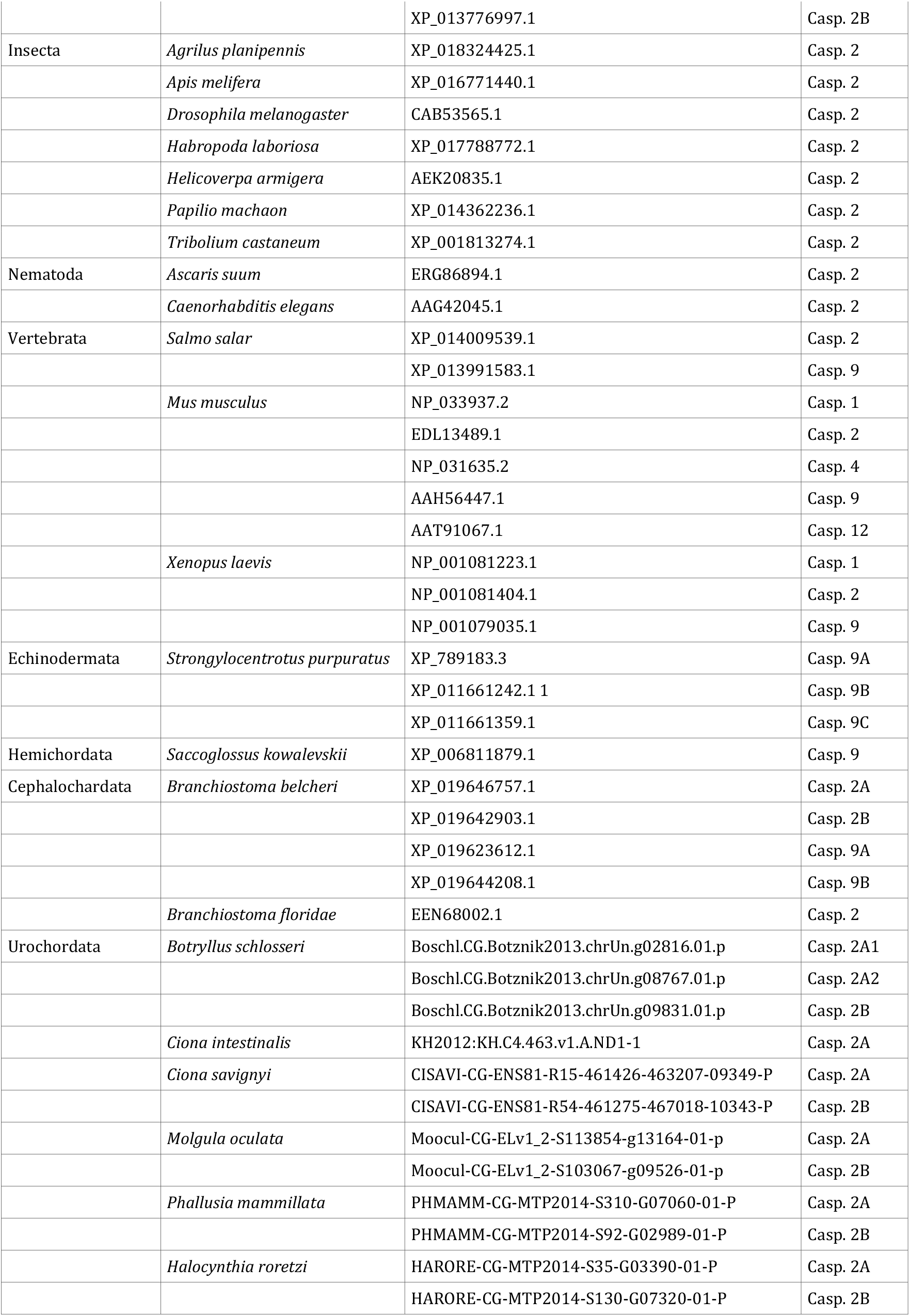
List of CARD-caspases used for phylogenetic analysis.

**Supplemental Table 2:** Topology tests between Bayesian inference method (BI topology) and maximum likelihood estimation (ML topology). p-KH : p-value of one sided Kishino-Hasegawa test. p-SH : p-value of Shimodaira-Hasegawa test. p-AU : p-value of approximately unbiased (AU) test. In all cases, BI topology is more likelihood than the ML topology.

|  | p-KH | p-SH | p-AU |
| --- | --- | --- | --- |
| BI topology | 0.714 | 1 | 0.702 |
| ML topology | 0.286 | 0.286 | 0.298 |

**Supplemental Table 3:**
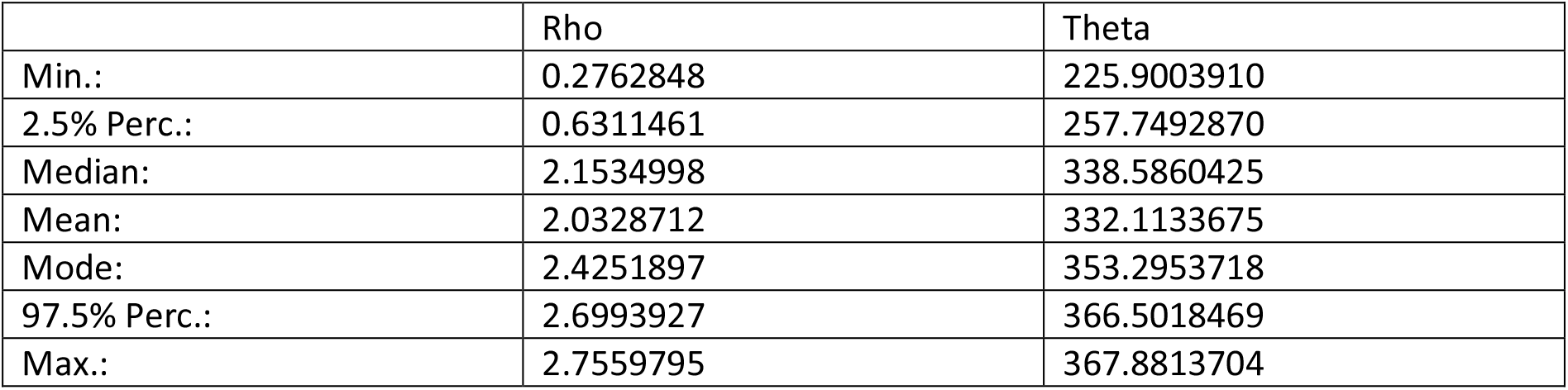
Estimation of the recombination (Rho) and substitution (Theta) rates using ProteinEvolverABC on metazoans CARD-caspases alignment. The low rho values indicate an absence of recombination and consequently no domain shuffling.

**Supplemental Table 4:**
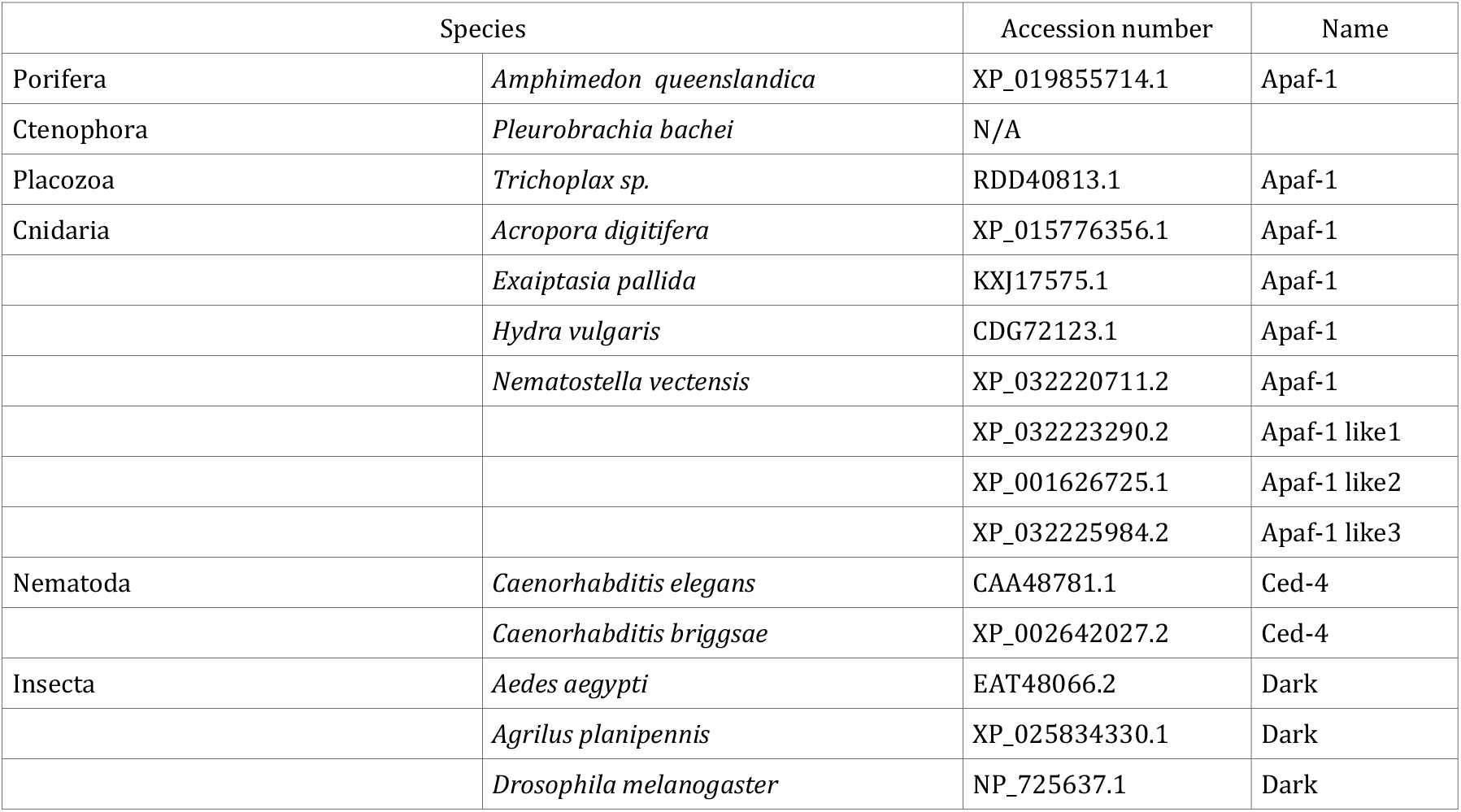
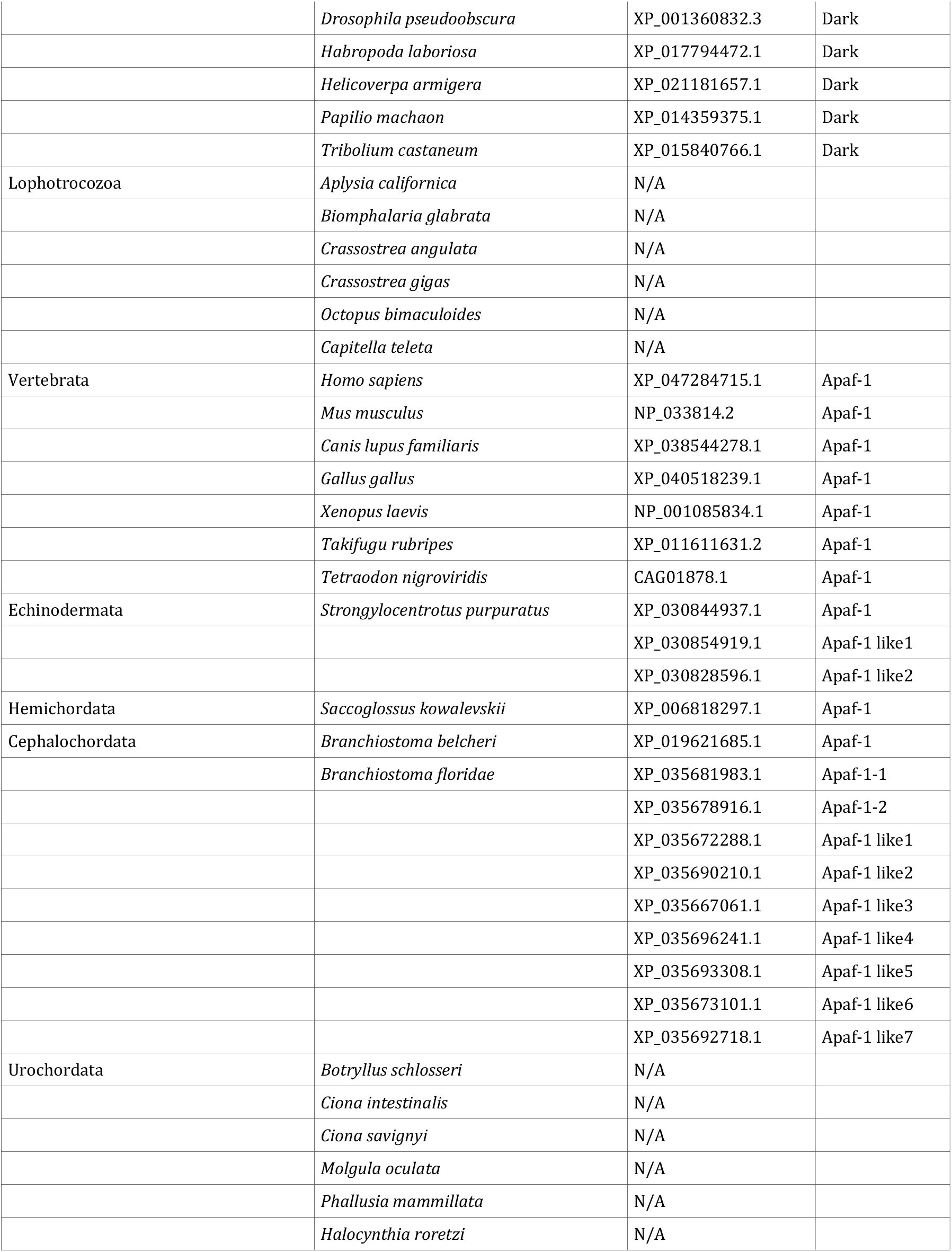
List of Apaf-1 among metazoans. N/A: No APAF-1 has been identified.

**Supplemental table 5:** List of Bcl-2 used for phylogenetic analysis.

| Species |  | Accession number | Names |
| --- | --- | --- | --- |
| Porifera | *Amphimedon queenslandica* | XP_003383425.1 | Bcl-2 like 1 |
|  |  | XP_003387574.1 | Bcl-2 like 2 |
| Cnidaria | *Hydra vulgaris* | EF104645.1 | Bak 1 |
|  |  | EU035760.1 | Bak 2 |
|  |  | XP_012562061.1 | Bax |
|  |  | EU035764.1 | Bcl-2 like 1 |
|  |  | EF104646.1 | Bcl-2 like 2 |
|  |  | EU035765.1 | Bcl-2 like 3 |
|  |  | EU035763.1 | Bcl-2 like 4 |
|  |  | EU035761.1 | Bok |
|  |  | EF104647.1 | Mcl-1 1 |
|  |  | EU035762 | Mcl-1 2 |
| Nematoda | *Caenorhabditis elegans* | AAA20080.1 | Ced-9 |
| Insecta | *Drosophila melanogaster* | AAF44120.1 | Buffy (Bok-1) |
|  |  | AAF26289.1 | Buffy (Bok-2) |
| Lophotrocozoa | *Biomphalaria glabrata* | XP_013085524.1 | Bax 1 |
|  |  | XP_013096338_1 | Bax 2 |
|  |  | XP_013086802.1 | Bax 3 |
|  |  | XP_013070177.1 | Bak |
|  |  | XP_013081872.1 | Bcl-2 like 1 |
|  |  | XP_013093137.1 | Bcl-2 like 2 |
|  |  | XP_013068612.1 | Bcl-2 like 3 |
|  |  | XP_013083436.1 | Bcl-2 like 4 |
|  |  | XP_013069706.1 | Bok |
|  |  | XP_013065969.1 | Mcl-1 |
|  | *Crassostrea gigas* | XP_011424481.1 | Bax |
|  |  | XP_011439700.1 | Bak |
|  |  | XP_011449013.1 | Bcl-2 like 1 |
|  |  | ACH42081.1 | Bcl-2 like 2 |
|  |  | EKC18663.1 | Bcl-2 like 3 |
|  |  | XP_011436990.1 | Mcl-1 1 |
|  |  | EKC40007.1 | Mcl-1 2 |
| Vertebrata | *Homo sapiens* | AAA74466.1 | Bak |
|  |  | NP_001278357.1 | Bax |
|  |  | API71171.1 | Bcl-2 |
|  |  | AAK48715.1 | Bcl-B |
|  |  | AAB09055.1 | Bcl-w |
|  |  | CAA80661.1 | Bcl-xL |
|  |  | NP_004040.1 | Bfl-1 |
|  |  | NP_115904.1 | Bok |
|  |  | AAF64255.1 | Mcl-1 |
|  | *Mus musculus* | NP_031549.2 | Bak |
|  |  | NP_031553.1 | Bax |
|  |  | AAH95964.1 | Bcl-2 |
|  |  | Q9Z0F3.1 | Bcl-B |
|  |  | AAA51039.1 | Bcl-xL |
|  |  | AAB09056.1 | Bcl-w |
|  |  | Q07440.1 | Bfl-1 |
|  |  | NP_058058.1 | Bok |
|  |  | NP_032588.1 | Mcl-1 |
|  | *Xenopus laevis* | NP_001089587.1 | Bak |
|  |  | AAR84081.1 | Bax |
|  |  | BAH28834.1 | Bcl-2 |
|  |  | AAI10791.1 | Bcl-xl |
|  |  | XP_018089640.1 | Bcl-w |
|  |  | NP_001139563.1 | Bok |
|  |  | ACI47310.1 | Mcl-1 |
| Urochordata | *Ciona intestinalis* | KH2012:KH.C4.794.v1.A.SL1-1 | Bax |
|  |  | KH2012:KH.S653.2.v2.A.SL1-1 | Bcl-xl |
|  |  | KH2012:KH.L87.39.v1.A.ND1-1 | Bok |
|  | *Molgula oculata* | Moocul.CG.ELv1_2.S96550.g08147.01.p | Bax |
|  |  | Moocul.CG.ELv1_2.S112899.g12639.01.p | Bok |

## BIBLIOGRAPHY

1. Jacobson, M. D., Weil, M. & Raff, M. C. Programmed cell death in animal development. Cell 88, 347–354 (1997).

2. Hipfner, D. R. & Cohen, S. M. Connecting proliferation and apoptosis in development and disease. Nat Rev Mol Cell Biol 5, 805–815 (2004).

3. Vriz, S., Reiter, S. & Galliot, B. Cell death: a program to regenerate. Curr. Top. Dev. Biol. 108, 121–151 (2014).

4. Hengartner, M. O. The biochemistry of apoptosis. Nature 407, 770–776 (2000).

5. Lettre, G. & Hengartner, M. O. Developmental apoptosis in C. elegans: a complex CEDnario. Nat Rev Mol Cell Biol 7, 97–108 (2006).

6. Ellis, H. M. & Horvitz, H. R. Genetic control of programmed cell death in the nematode C. elegans. Cell 44, 817–829 (1986).

7. Bender, C. E. et al. Mitochondrial pathway of apoptosis is ancestral in metazoans. Proc. Natl. Acad. Sci. U.S.A. 109, 4904–4909 (2012).

8. Steller, H. Regulation of apoptosis in Drosophila. Cell Death Differ. 15, 1132–1138 (2008).

9. Driscoll, M. Cell death in C. elegans: molecular insights into mechanisms conserved between nematodes and mammals. Brain Pathol 6, 411–425 (1996).

10. Dorstyn, L., Akey, C. W. & Kumar, S. New insights into apoptosome structure and function. Cell Death Differ 25, 1194–1208 (2018).

11. Young, N. D. et al. Diversity in the intrinsic apoptosis pathway of nematodes. Commun Biol 3, 478 (2020).

12. Bell, R. A. V. & Megeney, L. A. Evolution of caspase-mediated cell death and differentiation: twins separated at birth. Cell Death Differ 24, 1359–1368 (2017).

13. Shrestha, S. et al. Caspases from scleractinian coral show unique regulatory features. J Biol Chem 295, 14578–14591 (2020).

14. Uren, A. G. et al. Identification of Paracaspases and Metacaspases: Two Ancient Families of Caspase-like Proteins, One of which Plays a Key Role in MALT Lymphoma. Molecular Cell 6, 961–967 (2000).

15. Crawford, E. D. et al. Conservation of caspase substrates across metazoans suggests hierarchical importance of signaling pathways over specific targets and cleavage site motifs in apoptosis. Cell Death Differ 19, 2040–2048 (2012).

16. Thornberry, N. A. & Lazebnik, Y. Caspases: enemies within. Science 281, 1312–1316 (1998).

17. S, E. Apoptosis: a review of programmed cell death. Toxicologic pathology vol. 35 https://pubmed.ncbi.nlm.nih.gov/17562483/ (2007).

18. Galluzzi, L. et al. Molecular mechanisms of cell death: recommendations of the Nomenclature Committee on Cell Death 2018. Cell Death Differ. 25, 486–541 (2018).

19. Terajima, D. et al. Identification of candidate genes encoding the core components of the cell death machinery in the Ciona intestinalis genome. Cell Death Differ. 10, 749–753 (2003).

20. Delsuc, F., Brinkmann, H., Chourrout, D. & Philippe, H. Tunicates and not cephalochordates are the closest living relatives of vertebrates. Nature 439, 965–968 (2006).

21. Lee, T. V., Kamber Kaya, H. E., Simin, R., Baehrecke, E. H. & Bergmann, A. The initiator caspase Dronc is subject of enhanced autophagy upon proteasome impairment in Drosophila. Cell Death Differ 23, 1555–1564 (2016).

22. Fogarty, C. E. et al. Extracellular Reactive Oxygen Species Drive Apoptosis-Induced Proliferation via Drosophila Macrophages. Curr Biol 26, 575–584 (2016).

23. Fan, Y. et al. Genetic models of apoptosis-induced proliferation decipher activation of JNK and identify a requirement of EGFR signaling for tissue regenerative responses in Drosophila. PLoS Genet 10, e1004131 (2014).

24. Fan, Y. & Bergmann, A. The cleaved-Caspase-3 antibody is a marker of Caspase-9-like DRONC activity in Drosophila. Cell Death Differ 17, 534–539 (2010).

25. Napoletano, F. et al. p53-dependent programmed necrosis controls germ cell homeostasis during spermatogenesis. PLoS Genet 13, e1007024 (2017).

26. Lim, Y., Dorstyn, L. & Kumar, S. The p53-caspase-2 axis in the cell cycle and DNA damage response. Exp Mol Med 53, 517–527 (2021).

27. Bao, Q. & Shi, Y. Apoptosome: a platform for the activation of initiator caspases. Cell Death Differ 14, 56–65 (2007).

28. Jaroszewski, L., Rychlewski, L., Reed, J. C. & Godzik, A. ATP-activated oligomerization as a mechanism for apoptosis regulation: fold and mechanism prediction for CED-4. Proteins 39, 197–203 (2000).

29. Arenas, M. ProteinEvolverABC: Coestimation of Recombination and Substitution Rates in Protein Sequences by approximate Bayesian computation. Bioinformatics btab617 (2021) doi:10.1093/bioinformatics/btab617.

30. Lulla, V. & Firth, A. E. A hidden gene in astroviruses encodes a viroporin. Nat Commun 11, 4070 (2020).

31. Nieva, J. L., Madan, V. & Carrasco, L. Viroporins: structure and biological functions. Nat Rev Microbiol 10, 563–574 (2012).

32. Zmasek, C. M. & Godzik, A. Evolution of the animal apoptosis network. Cold Spring Harb Perspect Biol 5, a008649 (2013).

33. Zmasek, C. M., Zhang, Q., Ye, Y. & Godzik, A. Surprising complexity of the ancestral apoptosis network. Genome Biol. 8, R226 (2007).

34. Zou, H., Henzel, W. J., Liu, X., Lutschg, A. & Wang, X. Apaf-1, a human protein homologous to C. elegans CED-4, participates in cytochrome c-dependent activation of caspase-3. Cell 90, 405–413 (1997).

35. Zou, H., Li, Y., Liu, X. & Wang, X. An APAF-1.cytochrome c multimeric complex is a functional apoptosome that activates procaspase-9. J. Biol. Chem. 274, 11549–11556 (1999).

36. Dorstyn, L. et al. The role of cytochrome c in caspase activation in Drosophila melanogaster cells. J Cell Biol 156, 1089–1098 (2002).

37. Shi, Y. Mechanical aspects of apoptosome assembly. Curr Opin Cell Biol 18, 677–684 (2006).

38. Li, Y. et al. Conservation and divergence of mitochondrial apoptosis pathway in the Pacific oyster, Crassostrea gigas. Cell Death Dis 8, e2915 (2017).

39. Kiss, T. Apoptosis and its functional significance in molluscs. Apoptosis 15, 313–321 (2010).

40. Romero, A., Novoa, B. & Figueras, A. The complexity of apoptotic cell death in mollusks: An update. Fish Shellfish Immunol. 46, 79–87 (2015).

41. Estévez-Calvar, N., Romero, A., Figueras, A. & Novoa, B. Genes of the mitochondrial apoptotic pathway in Mytilus galloprovincialis. PLoS ONE 8, e61502 (2013).

42. Srivastava, M. et al. The Amphimedon queenslandica genome and the evolution of animal complexity. Nature 466, 720–726 (2010).

43. Srivastava, M. et al. The Trichoplax genome and the nature of placozoans. Nature 454, 955–960 (2008).

44. Zermati, Y. et al. Nonapoptotic role for Apaf-1 in the DNA damage checkpoint. Mol Cell 28, 624–637 (2007).

45. Ferraro, E. et al. Apaf1 plays a pro-survival role by regulating centrosome morphology and function. J Cell Sci 124, 3450–3463 (2011).

46. Shakeri, R., Kheirollahi, A. & Davoodi, J. Apaf-1: Regulation and function in cell death. Biochimie 135, 111–125 (2017).

47. Lee, E. F. et al. Discovery and molecular characterization of a Bcl-2-regulated cell death pathway in schistosomes. Proc. Natl. Acad. Sci. U.S.A. 108, 6999–7003 (2011).

48. Banjara, S., Suraweera, C. D., Hinds, M. G. & Kvansakul, M. The Bcl-2 Family: Ancient Origins, Conserved Structures, and Divergent Mechanisms. Biomolecules 10, (2020).

49. Kalkavan, H. & Green, D. R. MOMP, cell suicide as a BCL-2 family business. Cell Death Differ 25, 46–55 (2018).

50. Dunn, S. R., Phillips, W. S., Spatafora, J. W., Green, D. R. & Weis, V. M. Highly conserved caspase and Bcl-2 homologues from the sea anemone Aiptasia pallida: lower metazoans as models for the study of apoptosis evolution. J. Mol. Evol. 63, 95–107 (2006).

51. Clavier, A., Rincheval-Arnold, A., Colin, J., Mignotte, B. & Guénal, I. Apoptosis in Drosophila: which role for mitochondria? Apoptosis 21, 239–251 (2016).

52. Krumschnabel, G., Sohm, B., Bock, F., Manzl, C. & Villunger, A. The enigma of caspase-2: the laymen’s view. Cell Death & Differentiation 16, 195–207 (2009).

53. Olsson, M., Forsberg, J. & Zhivotovsky, B. Caspase-2: the reinvented enzyme. Oncogene 34, 1877–1882 (2015).

54. Lassus, P., Opitz-Araya, X. & Lazebnik, Y. Requirement for caspase-2 in stress-induced apoptosis before mitochondrial permeabilization. Science 297, 1352–1354 (2002).

55. Zhivotovsky, B. & Orrenius, S. Caspase-2 function in response to DNA damage. Biochem. Biophys. Res. Commun. 331, 859–867 (2005).

56. Krumschnabel, G., Manzl, C. & Villunger, A. Caspase-2: killer, savior and safeguard--emerging versatile roles for an ill-defined caspase. Oncogene 28, 3093–3096 (2009).

57. Olsson, M. et al. DISC-mediated activation of caspase-2 in DNA damage-induced apoptosis. Oncogene 28, 1949–1959 (2009).

58. Lavrik, I. N., Golks, A., Baumann, S. & Krammer, P. H. Caspase-2 is activated at the CD95 death-inducing signaling complex in the course of CD95-induced apoptosis. Blood 108, 559–565 (2006).

59. Braga, M. et al. Involvement of oxidative stress and caspase 2-mediated intrinsic pathway signaling in age-related increase in muscle cell apoptosis in mice. Apoptosis 13, 822–832 (2008).

60. Duan, H. & Dixit, V. M. RAIDD is a new ‘death’ adaptor molecule. Nature 385, 86–89 (1997).

61. Tinel, A. & Tschopp, J. The PIDDosome, a protein complex implicated in activation of caspase-2 in response to genotoxic stress. Science 304, 843–846 (2004).

62. Haupt, S., Berger, M., Goldberg, Z. & Haupt, Y. Apoptosis - the p53 network. J Cell Sci 116, 4077–4085 (2003).

63. Eyre-Walker, A. The genomic rate of adaptive evolution. Trends Ecol Evol 21, 569–575 (2006).

64. Gao, F. et al. EasyCodeML: A visual tool for analysis of selection using CodeML. Ecol Evol 9, 3891–3898 (2019).

65. Force, A. et al. Preservation of duplicate genes by complementary, degenerative mutations. Genetics 151, 1531–1545 (1999).

66. Conant, G. C. & Wolfe, K. H. Turning a hobby into a job: how duplicated genes find new functions. Nat Rev Genet 9, 938–950 (2008).

67. Fares, M. A., Keane, O. M., Toft, C., Carretero-Paulet, L. & Jones, G. W. The roles of whole-genome and small-scale duplications in the functional specialization of Saccharomyces cerevisiae genes. PLoS Genet 9, e1003176 (2013).

68. Meier, P., Finch, A. & Evan, G. Apoptosis in development. Nature 407, 796–801 (2000).

69. Degterev, A., Boyce, M. & Yuan, J. A decade of caspases. Oncogene 22, 8543–8567 (2003).

70. Fuchs, Y. & Steller, H. Live to die another way: modes of programmed cell death and the signals emanating from dying cells. Nat. Rev. Mol. Cell Biol. 16, 329–344 (2015).

71. Mancini, M. et al. Caspase-2 Is Localized at the Golgi Complex and Cleaves Golgin-160 during Apoptosis. J Cell Biol 149, 603–612 (2000).

72. Ouyang, Y. et al. Dronc caspase exerts a non-apoptotic function to restrain phospho-Numb-induced ectopic neuroblast formation in Drosophila. Development 138, 2185–2196 (2011).

73. Huh, J. R., Guo, M. & Hay, B. A. Compensatory proliferation induced by cell death in the Drosophila wing disc requires activity of the apical cell death caspase Dronc in a nonapoptotic role. Curr. Biol. 14, 1262–1266 (2004).

74. Kumar, S. & Doumanis, J. The fly caspases. Cell Death Differ 7, 1039–1044 (2000).

75. Vogeler, S., Carboni, S., Li, X. & Joyce, A. Phylogenetic analysis of the caspase family in bivalves: implications for programmed cell death, immune response and development. BMC Genomics 22, 80 (2021).

76. Pirger, Z., Rácz, B. & Kiss, T. Dopamine-induced programmed cell death is associated with cytochrome c release and caspase-3 activation in snail salivary gland cells. Biol. Cell 101, 105–116 (2009).

77. Romero, A., Estévez-Calvar, N., Dios, S., Figueras, A. & Novoa, B. New insights into the apoptotic process in mollusks: characterization of caspase genes in Mytilus galloprovincialis. PLoS ONE 6, e17003 (2011).

78. Zhang, L., Li, L. & Zhang, G. Gene discovery, comparative analysis and expression profile reveal the complexity of the Crassostrea gigas apoptosis system. Dev. Comp. Immunol. 35, 603–610 (2011).

79. Yang, B. et al. Molecular cloning of two molluscan caspases and gene functional analysis during Crassostrea angulata (Fujian oyster) larval metamorphosis. Mol. Biol. Rep. 42, 963–975 (2015).

80. Plachetzki, D. C., Pankey, M. S., MacManes, M. D., Lesser, M. P. & Walker, C. W. The Genome of the Softshell Clam Mya arenaria and the Evolution of Apoptosis. Genome Biol Evol 12, 1681–1693 (2020).

81. Piquet, B., Shillito, B., Lallier, F. H., Duperron, S. & Andersen, A. C. High rates of apoptosis visualized in the symbiont-bearing gills of deep-sea Bathymodiolus mussels. PLoS ONE 14, e0211499 (2019).

82. Sokolova, I. M., Evans, S. & Hughes, F. M. Cadmium-induced apoptosis in oyster hemocytes involves disturbance of cellular energy balance but no mitochondrial permeability transition. J. Exp. Biol. 207, 3369–3380 (2004).

83. Sokolova, I. M. Apoptosis in molluscan immune defense. Invertebrate Survival Journal 6, 49–58 (2009).

84. An, H.-K. et al. CASP9 (caspase 9) is essential for autophagosome maturation through regulation of mitochondrial homeostasis. Autophagy 16, 1598–1617 (2020).

85. Madadi, Z., Akbari-Birgani, S., Monfared, P. D. & Mohammadi, S. The non-apoptotic role of caspase-9 promotes differentiation in leukemic cells. Biochim Biophys Acta Mol Cell Res 1866, 118524 (2019).

86. Hollville, E. & Deshmukh, M. Physiological functions of non-apoptotic caspase activity in the nervous system. Semin Cell Dev Biol 82, 127–136 (2018).

87. Tran, H. T. et al. Caspase-9 has a nonapoptotic function in Xenopus embryonic primitive blood formation. J Cell Sci 130, 2371–2381 (2017).

88. Tamura, R. et al. Starfish Apaf-1 activates effector caspase-3/9 upon apoptosis of aged eggs. Sci Rep 8, 1611 (2018).

89. Belyi, V. A. et al. The origins and evolution of the p53 family of genes. Cold Spring Harb Perspect Biol 2, a001198 (2010).

90. Lu, W.-J., Amatruda, J. F. & Abrams, J. M. p53 ancestry: gazing through an evolutionary lens. Nat Rev Cancer 9, 758–762 (2009).

91. Berthelet, J. & Dubrez, L. Regulation of Apoptosis by Inhibitors of Apoptosis (IAPs). Cells 2, 163–187 (2013).

92. Ribeiro Lopes, M., Parisot, N., Callaerts, P. & Calevro, F. Genetic Diversity of the Apoptotic Pathway in Insects. in Evolution, Origin of Life, Concepts and Methods (ed. Pontarotti, P.) 253–285 (Springer International Publishing, 2019). doi:10.1007/978-3-030-30363-1_13.

93. Ribeiro Lopes, M. et al. Evolutionary novelty in the apoptotic pathway of aphids. Proc Natl Acad Sci U S A 117, 32545–32556 (2020).

94. Aubrey, B. J. et al. Mutant TRP53 exerts a target gene-selective dominant-negative effect to drive tumor development. Genes Dev 32, 1420–1429 (2018).

95. Aubrey, B. J., Kelly, G. L., Janic, A., Herold, M. J. & Strasser, A. How does p53 induce apoptosis and how does this relate to p53-mediated tumour suppression? Cell Death Differ 25, 104–113 (2018).

96. Gattiker, A., Gasteiger, E. & Bairoch, A. ScanProsite: a reference implementation of a PROSITE scanning tool. Appl Bioinformatics 1, 107–108 (2002).

97. Quevillon, E. et al. InterProScan: protein domains identifier. Nucleic Acids Res 33, W116–120 (2005).

98. Katoh, K. & Standley, D. M. MAFFT multiple sequence alignment software version 7: improvements in performance and usability. Mol. Biol. Evol. 30, 772–780 (2013).

99. Sievers, F. et al. Fast, scalable generation of high-quality protein multiple sequence alignments using Clustal Omega. Mol. Syst. Biol. 7, 539 (2011).

100. Hall, T. A. BioEdit : a user-friendly biological sequence alignment editor and analysis program for Windows 95/98/NT. Nucleic Acids Symp. Ser. 41, 95–98 (1999).

101. Castresana, J. Selection of conserved blocks from multiple alignments for their use in phylogenetic analysis. Mol. Biol. Evol. 17, 540–552 (2000).

102. Guindon, S. et al. New algorithms and methods to estimate maximum-likelihood phylogenies: assessing the performance of PhyML 3.0. Syst. Biol. 59, 307–321 (2010).

103. Gouy, M., Guindon, S. & Gascuel, O. SeaView version 4: A multiplatform graphical user interface for sequence alignment and phylogenetic tree building. Mol Biol Evol 27, 221–224 (2010).

104. Tamura, K., Stecher, G. & Kumar, S. MEGA11: Molecular Evolutionary Genetics Analysis Version 11. Mol Biol Evol 38, 3022–3027 (2021).

105. Ronquist, F. & Huelsenbeck, J. P. MrBayes 3: Bayesian phylogenetic inference under mixed models. Bioinformatics 19, 1572–1574 (2003).

106. Nguyen, L.-T., Schmidt, H. A., von Haeseler, A. & Minh, B. Q. IQ-TREE: A Fast and Effective Stochastic Algorithm for Estimating Maximum-Likelihood Phylogenies. Mol Biol Evol 32, 268–274 (2015).

107. Kishino, H. & Hasegawa, M. Evaluation of the maximum likelihood estimate of the evolutionary tree topologies from DNA sequence data, and the branching order in hominoidea. J Mol Evol 29, 170–179 (1989).

108. Shimodaira, H. An approximately unbiased test of phylogenetic tree selection. Syst Biol 51, 492–508 (2002).

109. Suyama, M., Torrents, D. & Bork, P. PAL2NAL: robust conversion of protein sequence alignments into the corresponding codon alignments. Nucleic Acids Res 34, W609–612 (2006).

